# Spatial landmark detection and tissue registration with deep learning

**DOI:** 10.1101/2023.08.24.554614

**Authors:** Markus Ekvall, Ludvig Bergenstråhle, Alma Andersson, Paulo Czarnewski, Johannes Olegård, Lukas Käll, Joakim Lundeberg

## Abstract

Spatial landmarks are crucial in describing histological features between samples or sites, tracking regions of interest in microscopy, and registering tissue samples within a common coordinate framework. Although other studies have explored unsupervised landmark detection, existing methods are not well-suited for histological image data as they often require a large number of images to converge, are unable to handle non-linear deformations between tissue sections, and are ineffective for z-stack alignment, other modalities beyond image data, or multimodal data. We address these challenges by introducing a new landmark detection and registration method, utilizing neural-network-guided thin-plate splines. Our proposed method is evaluated on a diverse range of datasets, demonstrating superior performance in both accuracy and stability compared to existing approaches.

## Main

Spatial landmarks are helpful in various areas of biotechnology. For instance, they are valuable when comparing histological heterogeneity between sites or samples^1^, keeping track of regions of interest in microscopy^2^, or registering tissue samples and transferring them to a common coordinate framework^3^. One can obtain spatial landmarks with, for example, experimental labeling^4^, microscopy^2^ techniques, and software-based manual or semi-manual^5^ annotation. However, the labor-intensive nature of locating spatial landmarks presents a bottleneck in spatial omics data analysis. Automating this process could significantly boost the scalability of sizeable spatial omics experiments while obviating the reliance on manually curated annotations.

Researchers have explored automating spatial landmark detection using deep learning techniques in computer vision, with successful results in both supervised^6^ ^7^ and unsupervised settings ^8 9^. Unsupervised methods hold greater promise, as they can address the general shortage of labeled landmark datasets, particularly in the diverse field of spatial omics. These unsupervised algorithms typically consist of a landmark detector network that identifies landmarks in images and a generative model that utilizes landmarks to guide image registration.

While these models have shown promise in specific tasks and biological applications^10^, their broad adaptation for tissue-related datasets necessitates overcoming three main challenges:

1. Limited Datasets: Deep learning techniques often necessitate vast datasets, sometimes in the order of 100,000 training examples, to discern general patterns and avoid overfitting. However, multi-omics studies often include fewer than ten training samples.
2. Non-linear Transformations: Current methods predominantly focus on datasets involving more straightforward affine transformations, such as rotation, scaling, and translation. However, researchers often encounter images that require a combination of elastic and rigid transformations to integrate multiple images in biological contexts^11^.
3. Multi-Modal Data Handling: The methods must handle data from different modalities, such as histology stains, spatially resolved transcriptomics, and mass spectrometry imaging (MSI), and process these modalities concurrently.

Building on the work of Sanchez et al.^8^ we introduce ELD (Effortless Landmark Detection) to address these challenges. In this study, we highlight the performance of ELD across a range of applications, including single-modality data registration, 3D modeling, and multimodal data alignment. For single-modality data registration, we demonstrate ELD’s enhanced stability and efficiency across modalities such as Visium, H&E, and ISS. We also show that it outperforms other landmark detection models in numerous tests. Regarding 3D modeling, ELD’s proficiency was underscored by its notable improvement in registration metrics compared to eight other registration models on a mouse prostate dataset. Finally, we show that ELD can successfully model Visium and H&E or Visium and MSI data simultaneously, demonstrating its ability to learn modality-agnostic landmarks for integrating multimodal datasets.

## Results

### Benchmarking ELD against existing methods

One can design a deep neural network to better generalize for small datasets by adding constraints to the model, such as reducing the neuron count per layer, decreasing the number of layers, implementing dropout, or adding a regularizer to the loss function^10^. In ELD, we constrain the solution space by removing the generative network while retaining the landmark-detecting network, as suggested by Sanchez et al.^8^. With the image landmarks identified, registration can be easily performed using landmark-based methods, such as homography^11^ or thin-plate spline (TPS)^12^. However, given the elastic nature of the transformations in our data, we here use TPS for registration purposes. These methods offer several advantages, including having fewer unlearnable parameters (hard constraints) and being more computationally efficient than large deep neural networks.

The process of aligning tissue slices to a common coordinate framework (CCF) can be outlined as follows (Figure 1a): To begin, the ELD system employs an unsupervised trained spatial landmark detection network to pinpoint landmarks on the desired tissue slices or manual annotations can be used. Once these critical points have been established across all slices, ELD utilizes landmark-centric alignment techniques, such as TPS or homography, to align the regions. As a final step, ELD projects all the aligned tissue regions onto a CCF, facilitating comparative studies across various slices.

**Figure 1:**
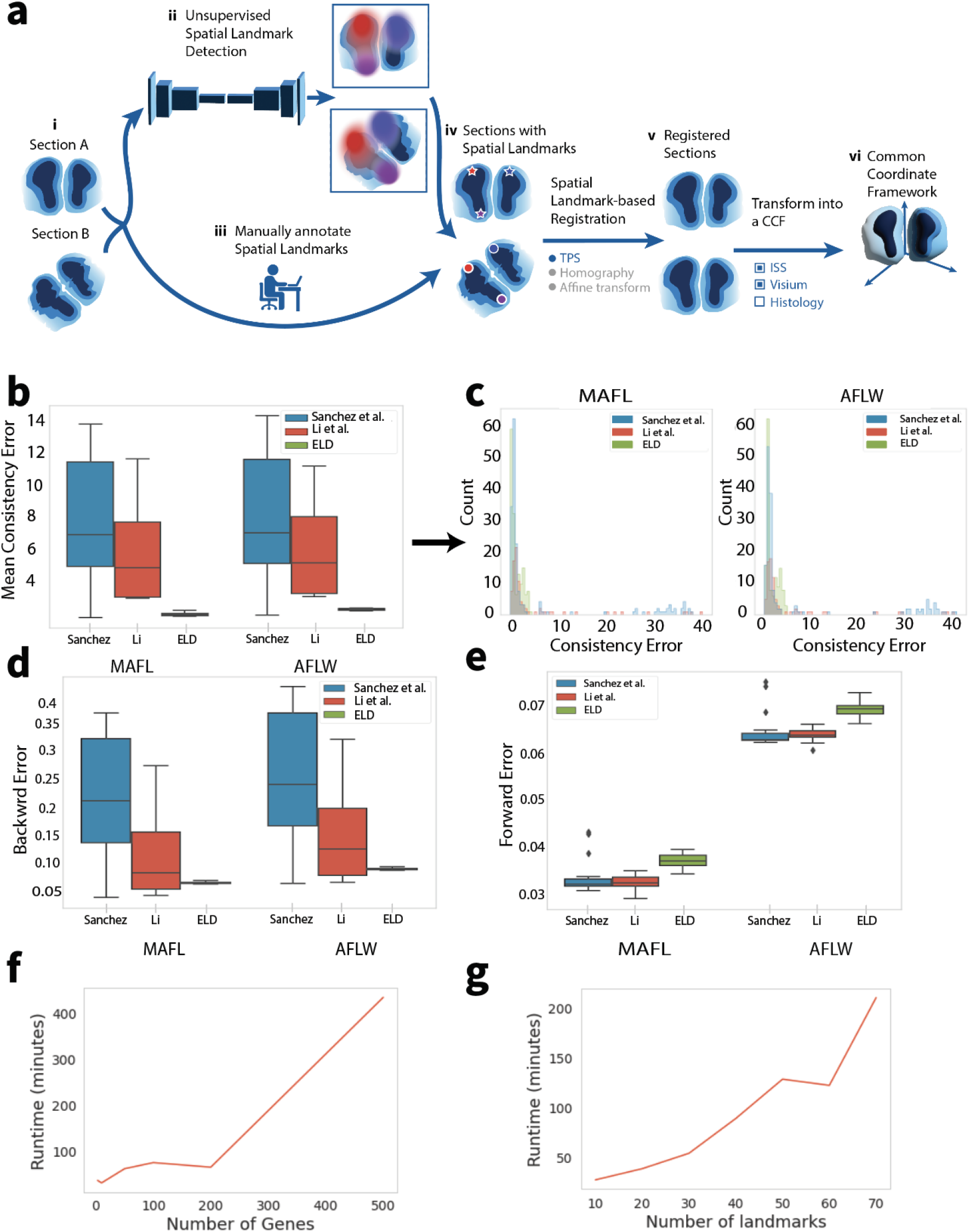
Overview of ELD framework and performance assessment. (a) Proposed workflow of acquiring spatial landmarks and aligning sections within a common coordinate framework. The process can be divided into the following steps: i) Obtain two different sections, A and B, for which we want to identify landmarks. ii) Use a trained unsupervised spatial landmark detection network or iii) manually annotate the sections to obtain landmarks for both sections. iv) After acquiring landmarks for both samples, register Section B to Section A using various landmark-based alignment methods, such as TPS or Homography. v) Obtain the registered sample, which aligns Section B with Section A. vi) Map the samples to a common coordinate framework, allowing cross-section comparisons and analysis. b-e, Performance benchmarks for ELD and two other state-of-the-art models for landmark detection. Benchmarks were conducted on the MAFL and AFLW datasets. Figures show the distribution of the mean consistency error for each image (b), consistency error for each landmark (c), and backward (d) and forward errors (e) for each image. f,g, Runtime analysis. Figures show the time required for convergence in minutes as a function of the number of genes or color channels used when the number of landmarks is set to 10 (f) and as a function of the number of landmarks used when the number of channels is set to 3 (g).

In this study, we employ standard error metrics to assess the performance of ELD and two other state-of-the-art landmark detection methods. These metrics include forward error, backward error, and consistency error^8^. Performance benchmarks are conducted using CelebA for training, and the MAFL and AFLW datasets for evaluation, which are frequently employed for tasks of this nature.

Consistency error evaluates landmark stability through geometric consistency. To calculate the consistency error, one must: 1) detect the landmarks in the image, apply an affine transformation to the landmarks, 2) apply the same affine transformation to the image, and then detect the landmarks again but on the transformed image. The error is determined by comparing the point-to-point distances between the two sets of landmarks. ELD exhibits superior consistency compared to other methods (Figure 1b). Interestingly, a closer examination of the results reveals that the performance difference is largely attributed to the other methods’ tendency to identify landmarks that are sometimes significantly misaligned (Figure 1c). While most landmarks are consistent, these outliers contribute to a much higher mean error than ELD.

The forward error is calculated by training a linear regression model using a set of manually annotated points and the detected landmarks. The trained regressor predicts the annotated points based on the detected landmarks. Conversely, the backward error is computed using a linear regression model trained in reverse order, i.e., using the annotated landmarks to predict the detected landmarks. This error serves as a measure of the stability of the detected landmarks. A model with a low forward error but a high backward error will likely detect a low number of stable landmarks. On the other hand, a model that has a low backward error but high forward error is likely to converge to a fixed set of points independent of the input image.

ELD exhibits significantly better backward error than other methods (Figure 1d), which can be attributed to the inconsistent landmarks found by the other methods. Although all models show better performance in forward error than backward error, ELD displays marginally worse performance in forward error (Figure 1e). This indicates that ELD sacrifices some generalization in favor of significantly improved consistency.

We conducted two tests to evaluate ELD’s runtime requirements: one with varying numbers of genes or image channels (Figure 1f) and another with varying numbers of landmarks (Figure 1g). As detailed in the methods section, the convergence criterion is quite stringent; however, convergence is typically achieved more quickly in real-world applications.

### Performance Evaluation on Single-modality Data

An effective registration method for Visium data, Eggplant^13^, is currently openly available. One limitation of Eggplant, however, is its reliance on manual spatial landmark annotation. Therefore, we next sought to test whether the spatial landmarks generated by ELD can supplant manual annotation and improve the performance of Eggplant. Using Eggplant to transfer the gene expression of three target genes with distinct expression patterns (*Nrgn, Apoe*, and *Omp*) in the mouse olfactory bulb to a reference section using either manually or automatically detected landmarks, we find that the landmarks produced by ELD yield results that are at least as accurate as those obtained using manual annotation (Figure 2a and 2b). For this experiment, we used 12 mouse olfactory bulb samples, the same reference as employed in Eggplant^13^. Our results are consistent whether landmarks were identified using histology or expression data from 3 or 100 genes.

**Figure 2:**
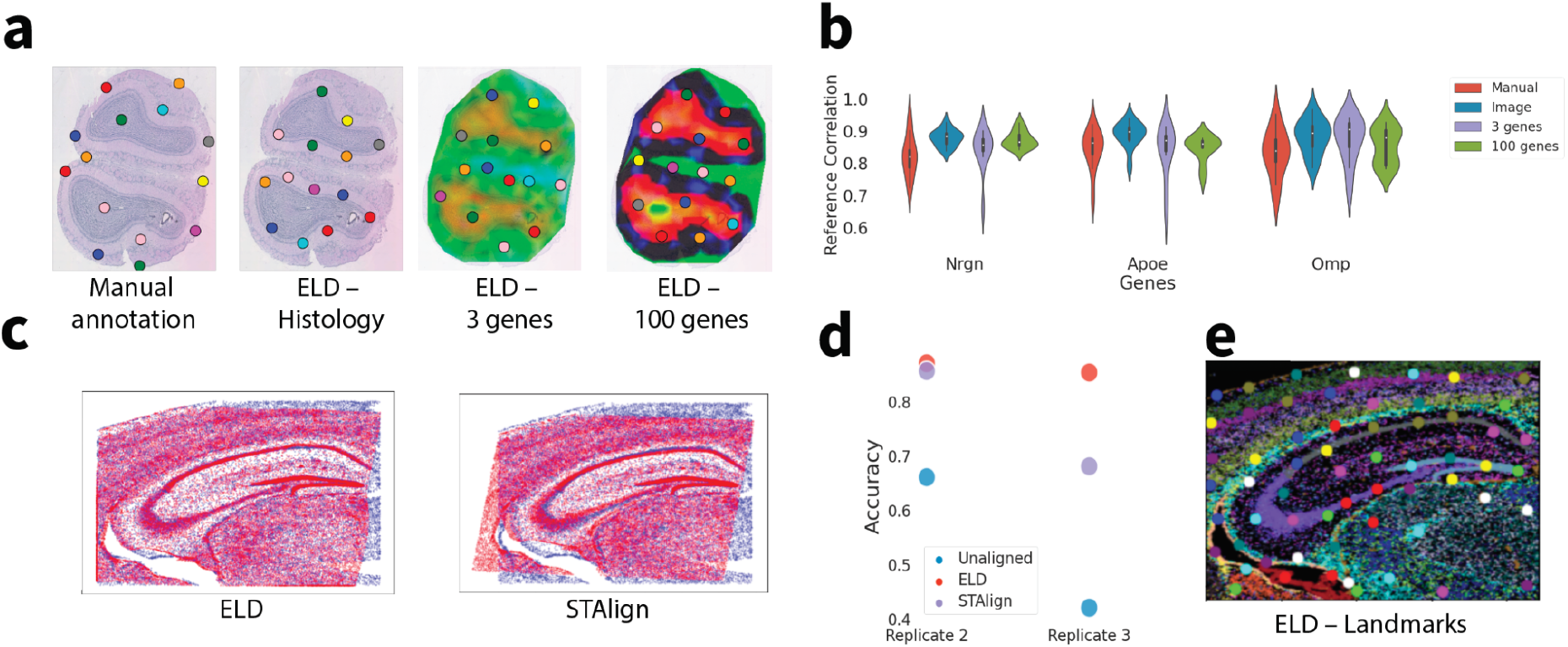
Performance on single-modality data. (a) Landmarks identified by ELD when trained on various modalities, such as histology and Visium data, alongside an image with manual annotation for comparison. The rightmost image, which includes 100 genes, is visualized using PCA; however, all genes were utilized during the training process. For all experiments, 14 landmarks were used. (b) Violin plots show the correlation of three target genes (*Nrgn, Apoe*, and *Omp*) between the ten registered samples and the reference. (c) Visual comparison of registration quality between ELD and STAlign for ISS data. (d) Evaluation of how accurately a k-nearest neighbor model predicts different anatomical regions. (e) ELD-predicted landmarks on an RGB image generated by clustering the ISS data.

We used three mouse brain coronal sections from S. Salas et al.^14^ to demonstrate ELD’s compatibility with ISS data. In this experiment, we employ RGB images of the clustering on the ISS data (Figure 2e) and use TPS for the final registration. To evaluate the effectiveness of the registration, we assess how well a simple k-nearest neighbor model trained on the reference could predict the correct anatomical region on the registered samples. Comparing ELD to STAlign^15^, which has shown promising results for aligning data from ISS experiments, we find that ELD attains a higher accuracy in both replicates (Figure 2d).

### 3D modeling

To make it possible to align a stack of multiple tissue sections, whose morphology may change drastically along the stacking axis, we modify ELD to generate anchor points instead of landmarks. The general procedure is illustrated in Figure 3a. Briefly, the most significant difference for z-stack alignment involves controlling how the area changes of the transformed tissue. This forces the landmarks to act more like anchor points with fixed x-y coordinates instead of identifying common morphology, as demonstrated in Figure 3c.

**Figure 3:**
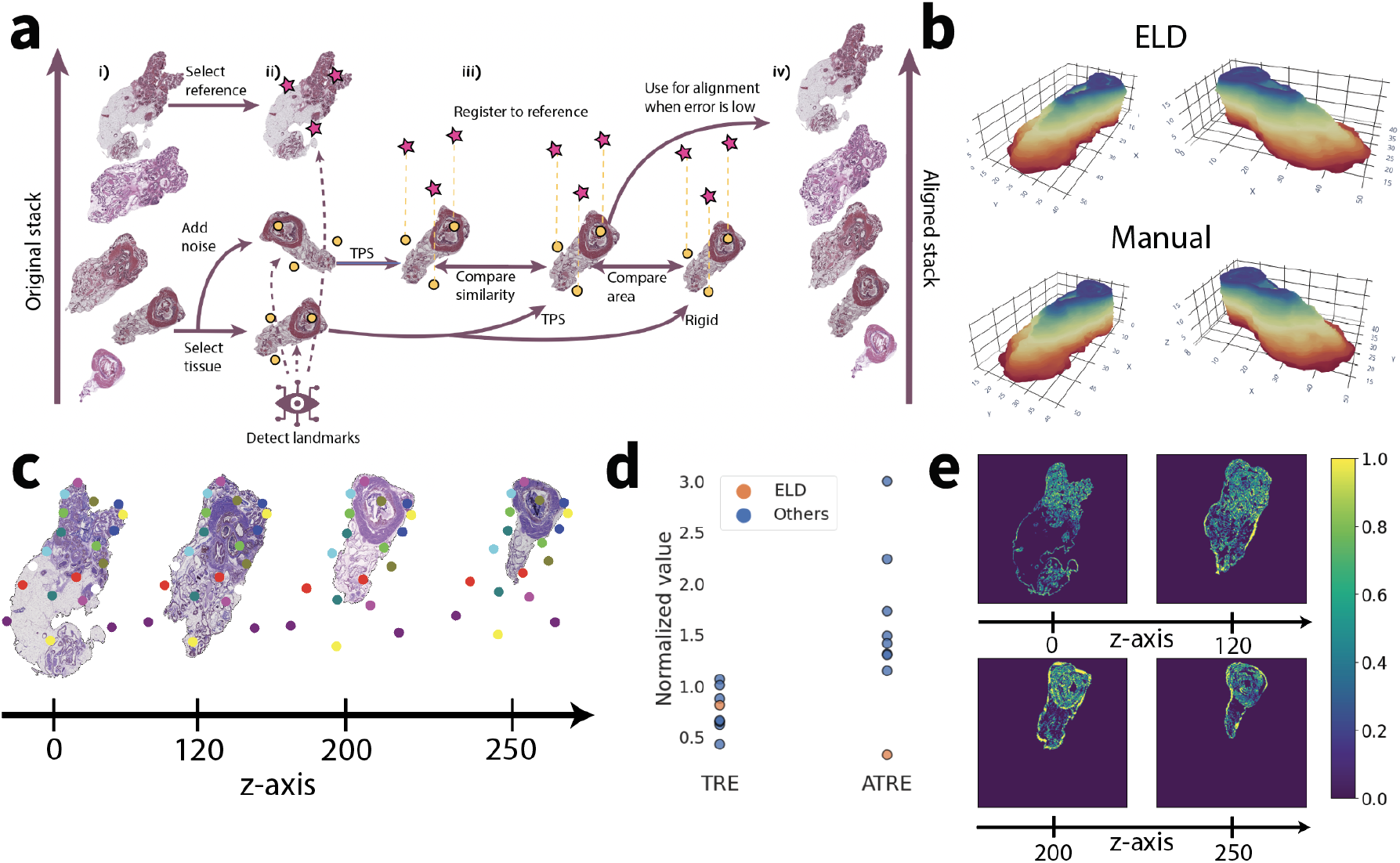
3D tissue reconstruction with ELD. (a) Illustration of the 3D modeling process. First, i) randomly select a reference from the tissue stack, choose a tissue, and create a noisy counterpart to map to the reference. Next, detect landmarks ii) for the reference and both the source tissue and its noisy counterpart. Then, map the noisy source image to the reference using TPS, and map the original source image with TPS and rigid transformation to the reference. Compute iii) the similarity loss between the mapped noisy source and the original source image, and compare the area change between the source image mapped with TPS and rigid transformation. Repeat this process for all tissues in the stack until convergence. Finally, map all tissues to the reference. (b) Demonstration of final registration using ELD, and with the manual annotations. (c) Display of aligned tissues with their anchors from the tissue stack. A total of 20 landmarks were used for the alignment. (d) Performance comparison of ELD and other models. All results are normalized using the value obtained when aligning with manual landmarks (corresponding to a score of 1). (e) Absolute error between manual alignment and alignment using ELD across four sections in the z-stack

To assess ELD’s 3D alignment performance, we utilize a mouse prostate dataset containing 260 slices from K. Kartasalo et al.^16^ The dataset contains annotations from two different annotators of four corresponding landmarks in each pair of consecutive sections. Additionally, we employ their published code to generate the results. Since most benchmarks were similar across all methods and different processing was done on the images, the root-mean-square error (RMSE) is difficult to compare fairly. Therefore, we chose to only present the landmark-related benchmarks Target to Registration Error (TRE) and Accumulated Target to Registration Error (ATRE). The TRE is calculated as the Euclidean distance between the actual and predicted locations of points. Specifically, these points are not used in the registration process (also known as target points), and this calculation is performed for each consecutive pair. The ATRE is a cumulative TRE over all the tissue sections. This measurement provides an overall indication of the total error in the registration task across all target points. The mean of TRE and ATRE is used, and it’s normalized by the score obtained when registering with manually annotated landmarks, as depicted in Figure 3d. While the performance of ELD is comparable to the other methods in terms of TRE, ELD significantly outperforms them in terms of ATRE, suggesting that the alignment is more consistent across the entire tissue volume. We compared eight other registration models, seven of which come from K. Kartasalo et al. ^16^ and CODA ^17^. The final 3D alignment is illustrated in Figure 3b.

### Performance Evaluation on Multimodal Data

ELD can detect landmarks and align tissue data from different modalities. To optimize the alignment between two distinct modalities, separate landmark detectors are used for each modality. During training, random samples from both modalities are selected, one sample is registered to the other, and their alignment is assessed in the latent space obtained from the landmark detector (Figure 4a).

**Figure 4:**
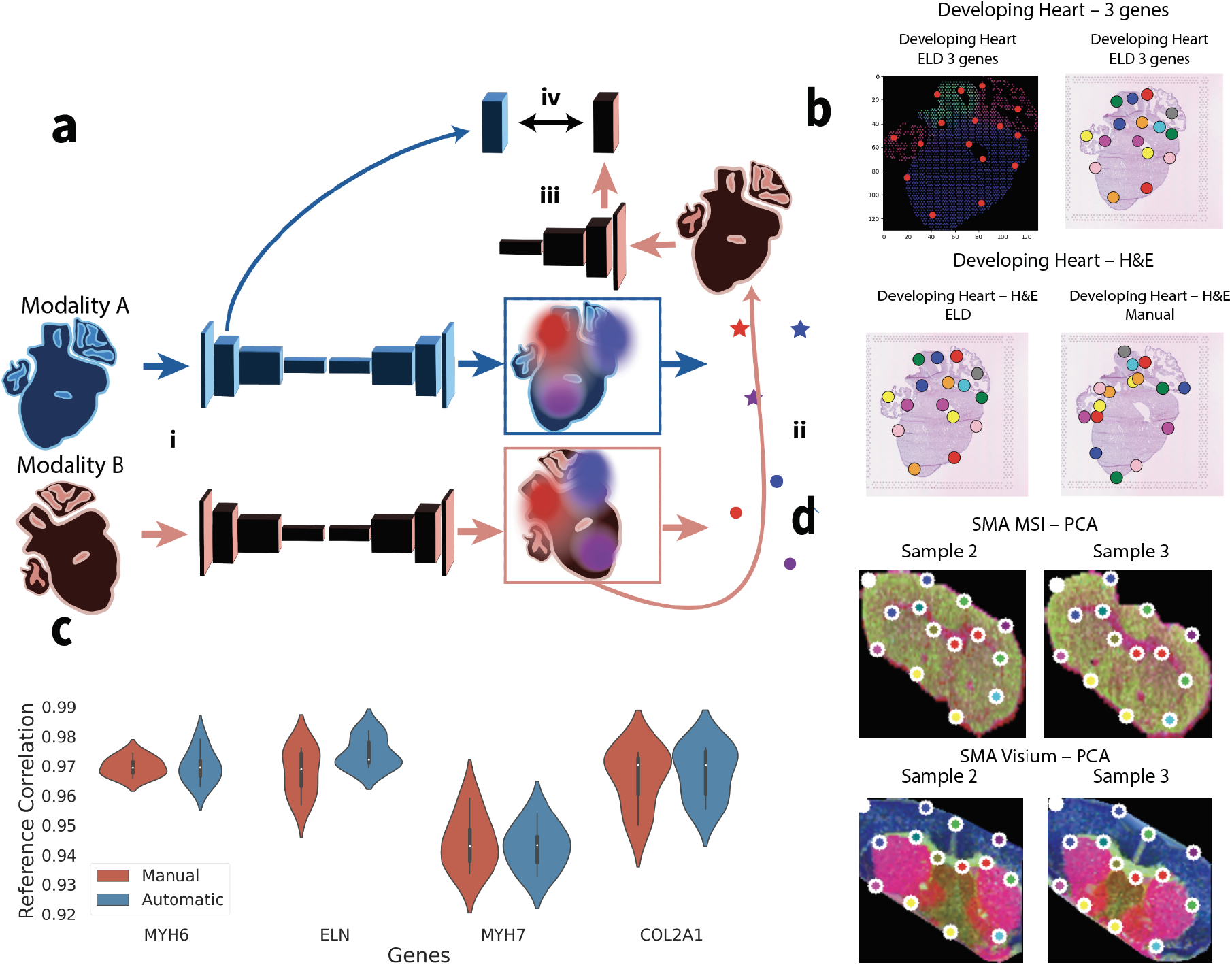
Benchmarking ELD for multimodal alignment. (a) Registration of multimodal data, overview. i) Each modality, A and B, is passed through its respective landmark detector, and a latent representation of modality A is saved. ii) Landmarks are extracted from the heatmaps, and the two modalities’ samples are registered. iii) A latent representation is obtained from modality B. iv) The similarity of the latent representations of modalities A and B is optimized. (b) Detected landmarks for gene expression and histology images. The upper-left image shows *MYH6, ELN*, and *MYH7 gene expression*. In total, 16 landmarks were used for this experiment. c) Performance comparison of Eggplant’s alignment using manually annotated landmarks and landmarks detected with ELD. d) PCA visualization of MSI and Visium data with their respective landmarks.

We used the Human Developing Heart dataset^13^, which consists of four samples, to demonstrate ELD’s ability to align tissues from two different modalities. Histology images were used for the first two samples, while the genes *MYH6, ELN*, and *MYH7* from Visium expression data were employed to construct an image for the other samples. The detected landmarks for the two modalities are displayed (Figure 4b).

To benchmark ELD’s performance, we randomly selected one of the samples as reference. Then we calculated the correlation of the source sample to the reference with Eggplant, using both ELD’s landmarks and manually annotated landmarks. The programmatically detected landmarks perform comparably to manually annotated landmarks (Figure 4b-c).

To further demonstrate the flexibility of ELD to model data of diverse modalities, we apply it to PCA embeddings of MSI and Visium data. This data was extracted from three mouse striatum samples as per the study conducted by M. Vicari et al.^18^. For each of these samples, we employed a combination of both MSI and Visium methodologies. We find the generated landmarks to be qualitatively consistent across sections and to mark out biologically relevant anatomical features (Figure 4d).

## Discussion

In this study, we have introduced ELD (Effortless Landmark Detection), a novel method for unsupervised spatial landmark detection and registration that addresses the challenges of small datasets and non-linear transformations typically found in spatial omics and histological image data. ELD employs neural-network-guided thin-plate splines and outperforms existing approaches in terms of both accuracy and stability.

By removing the generative network and retaining the landmark-detecting network, ELD effectively addresses the issue of overfitting in small datasets by removing the generative network and retaining the landmark-detecting network. We have demonstrated that ELD achieves superior consistency and backward error performance compared to competing methods while showing marginally worse performance in forward error. Our runtime tests indicate that ELD is computationally efficient, and we have empirically observed convergence to be even quicker in many real-world applications.

By tweaking the optimization function, we have demonstrated the effectiveness of ELD in a wide range of applications, such as single-modality data registration, 3D modeling, and multimodal data alignment. Regarding single-modality data registration, ELD outperformed existing methods like manual annotation with Eggplant and STAlign. For 3D modeling, ELD was adapted to produce anchor points rather than landmarks, leading to successful z-stack alignment. Moreover, ELD significantly improved the ATRE metric for the mouse prostate dataset compared to eight competing models.

Finally, we have shown that ELD can align tissues from diverse modalities by utilizing distinct landmark detectors for each modality and comparing the registration similarity in a latent tissue space.

The primary objective of ELD is landmark detection, while registration serves as an added benefit. In this regard, relatively simple registration models, such as TPS, have been utilized. We believe that ELD has the potential to improve other models with more advanced registration approaches, like STAlign and CODA, similar to how it enhances Eggplant, by supplying ELD’s landmarks as a ground truth during the training phase.

Overall, ELD demonstrates a notable improvement over existing unsupervised landmark detection and registration methods in spatial omics and histological image data. Its versatility in addressing different data types and modalities makes it a promising tool for researchers in spatial biology.

## Methods

### Hardware

We used an NVIDIA A100-SXM 81GB graphics card, and 12 AMD EPYC 7742 64-Core Processors for all model training.

### Cost function: Multiscale Structural Similarity (MS-SSIM)

In all the experiments detailed in the subsequent sections, we utilize a cost function rooted in the Multi-Scale Structural Similarity (MS-SSIM) method. This approach allows for a comprehensive assessment of image quality by considering image details across a range of resolutions. This method extends the single-scale SSIM index, which compares luminance, contrast, and structure of two aligned signals, such as image patches^19^. MS-SSIM has been very useful in our experiments since we have due to the presence of significant batch effects, and MS-SSIM have demonstrated more robustness than for instance Mean Squared Error in our experiments.

The MS-SSIM procedure involves an iterative process of applying a low-pass filter to the image and downsampling the filtered image. Each iteration defines a new scale, culminating in the highest scale. Contrast and structure comparisons are computed at every scale, while luminance comparison is reserved for the highest scale^19^.

The overall quality assessment in MS-SSIM combines these measurements from all scales, using adjustable parameters for accounting for the relative importance of each component at every scale. The method yields a detailed image quality map, with the mean MS-SSIM index offering an overall evaluation of image quality. For a comprehensive understanding of MS-SSIM, refer to the work of Wang et al. ^19^.

For the calculation of MS-SSIM, we employed the PyTorch Image Quality Assessment ([PIQA](https://pypi.org/project/piqa/)) package, utilizing its default parameters along with a window size of 5. This configuration was chosen based on our preliminary trials, which indicated its effectiveness in our context.

### Landmark Drop-Out

When training ELD landmarks, they can become trapped in a local minimum, which often results in many landmarks occupying similar positions. To counteract this, we employ a technique known as landmark drop-out. This process involves the probabilistic removal of detected landmarks, with each landmark having a probability p of being dropped out. Empirical observations have demonstrated that this method allows the landmarks to escape local minima more rapidly, leading to a more diverse and satisfactory distribution of landmarks in a shorter time. Throughout all our experiments, we utilized a drop-out probability of 10%. Empirically, this has proven to work well in practice.

### Cropping

During training, data augmentation can occasionally cause images to be cut-off or contain missing regions due to batch effects. This can complicate the process when registering two images, as the missing portions can confuse the model. To mitigate this issue, we perform a cropping procedure on the registered and mapped images based on the black-colored background, which is represented by a 0 value across all channels. Specifically, we identify the masks for all black channels in both images and then crop both images according to these masks. To ensure precision, we implement a threshold of 0.1. This means any pixels in which all channels fall below this threshold are considered black.

### Registration: Thin-Plate Splines (TPS)

In all the experiments outlined in the following sections, we employ TPS for image registration. TPS is a widely used method known for its capacity to effectively handle image registration and deformation, primarily by interpolating scattered data points. TPS creates a smooth and flexible mapping between two sets of landmarks while minimizing bending energy. More formally, Given a set of *n* source points *p* and their corresponding target points *q*_*i*_ in 2D space, the goal is to find a mapping *f*(*x, y*) that minimizes the following energy function:

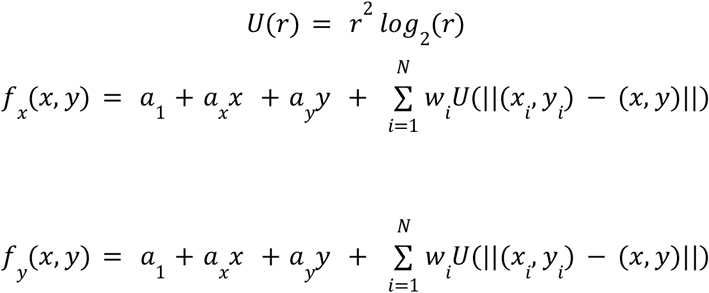

The TPS function, *f*(*x, y*), can be solved analytically to obtain the weights. Once these weights are acquired, the source image can be mapped to the reference. For a more in-depth understanding, we recommend referring to the study by Keller et al.^12^

TPS is of significant utility in scenarios demanding smooth and continuous transformations, such as shape morphing in computer graphics. Within the context of ELD training, TPS is leveraged to guide the learning of high-quality landmarks.

### Landmark Detector

The landmark detector employed in this article is identical to the one used by Sanchez et al. ^8^, which is an hourglass network consisting of approximately 6 million parameters.

### General image preprocessing

Throughout all experiments, we employed 128x128 images during training, achieved by utilizing cv2.resize and applying INTER_AREA-interpolation (referenced in docs.opencv.org) to transform the original image dimensions to the desired format. Furthermore, ELD cannot process flipped images, so it’s crucial to ensure all images are oriented in the same way prior to training.

### Data Augmentation

In every experiment, we utilized data augmentation strategies, including rotation, scaling, and elastic transformation. The rotation was implemented with a random angle selection between -15 and 15 degrees, paired with appropriate scaling to maintain the subject within the image frame. For the introduction of elastic noise, we employed the [elasticdeform](https://elasticdeform.readthedocs.io/en/latest/elasticdeform.html) package. Both the control points for the deformation grid and the sigma for the normal distribution were randomly selected within an experiment-dependent range.

### Visium preprocessing and filtering

For all Visium data, spots with fewer than 200 detected genes were removed, and genes present in less than three spots were also removed. The data were then normalized using [Scanpy](https://scanpy.readthedocs.io/en/stable/index.html) and log1p-transformed. The same genes as in Eggplant^13^ were chosen when selecting three genes. In the experiments where more genes were utilized, we performed Leiden clustering on neighborhood graphs derived from Principal Component Analysis (PCA). All methods were implemented with the default parameters provided by Scanpy. Subsequently, we employed Scanpy’s *rank_genes_groups* function, using the Leiden clusters as groupings and the t-test for ranking. This allowed us to select the top n-ranked genes per sample. Finally, the common genes across all samples were chosen for further analysis.

## Code availability

The code used in the experiments of this paper can be found at https://github.com/ekvall93/ELD.

## Data availability

All experimental data and models are available on Figshare at https://figshare.com/projects/ELD/167318. However, the 3D modeling data can be found separately on etsin.fairdata.fi

## Ethical declaration

The Regional Ethical Review Board in Stockholm and the National Board of Health and Welfare granted approval for the use of human developmental heart tissue in the study. The acquisition of the tissue, along with the data processing, complied with the ethical guidelines set forth by the Helsinki Convention (Dnr: 2:9/2015). The use of human foetal material from the elective routine abortions was approved by the Swedish National Board of Health and Welfare, and the analysis using this material was approved by the Swedish Ethical Review Authority (2018/769-31). The human developmental heart tissue, employed in this study, was obtained from medical abortions carried out at the Department of Gynecology, Danderyd Hospital, and Karolinska Huddinge Hospital. All the patients involved gave their informed consent in written form.

### Comparative Assessment of Landmark Quality and Runtime Requirement: Evaluating Existing Methods

During the training of ELD on the CelebA dataset for the purpose of landmark quality assessment, we create two augmented variants of each image: *X*_*target*_ and *X*_*source*_, as elaborated in the preceding section. *X*_*source*_ is then registered to *X*_*target*_ using the landmarks detected by the ELD, resulting in *X*_*registered*_. We then calculate the MS-SSIM loss between *X*_*registered*_ and *X*_*target*_, which is referred to as the base loss.

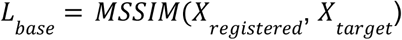

To guarantee consistency across images, we randomly select another sample, *Y*_*target*_, and align it to *X*_*target*_, forming *Y*_*registered*_. However, we only compute the MS-SSIM loss (with a window size of 3) between small patches surrounding the landmarks, specifically 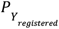 and 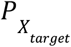. This is called the consistency loss and ensures that specific landmarks, such as the left eye, consistently target the same feature across different images.

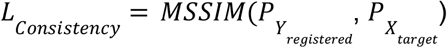

The primary loss is computed by combining the base loss and the consistency loss. The consistency loss is scaled by a factor of 0.1, determined through empirical testing, although a scalar in the range of 0.1 to 0.5 has been observed to yield similar performance. The final loss function is, therefore, a composite of these two components.

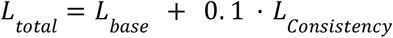

In this comparison, we evaluated our method against two other recently published landmark detection methods [9, 8], both of which represent the current state of the art in this field. We trained a total of 15 models for each method. We ran their models using default parameters for 80 epochs, a batch size of 48, and also trained our models with the same parameters, but added elastic noise with a sigma value of 3. The learning rate employed was 1e-4, using a learning rate scheduler with a step size of 10 epochs and a learning rate decay of 0.95. TPS was used to register the samples. All benchmarks were performed with the code from Sanchez et al.^8^.

In the runtime experiment, we used an initial learning rate of 1e-4 which was annealed by a learning rate scheduler with step size 3 and a learning rate decay of 0.95. TPS was employed for registration purposes. Samples were perturbed by elastic noise with a sigma parameter of 5.5. A batch size of 48 was used, resulting in 300 iterations per epoch. Training was stopped when the loss improved by less than 1e-4 over 10 consecutive epochs.

### Performance Evaluation on Single-modality Data

Maintaining the same objective outlined in the preceding section, but we modified the calculation of *L*_*Consistency*_. Instead of applying MS-SSIM to patches of the aligned sections, we computed it directly between *X*_*target*_ and *Y*_*registered*_. This adjustment is justifiable considering the minor batch effects present between samples. Consequently, the consistency loss, denoted as *L*_*Consistency*_, is calculated as the MS-SSIM between *Y*_*registered*_ and *X*_*target*_ :

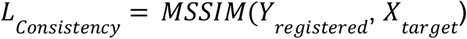

The final loss, *L*_*total*_, is then computed by combining the base loss, *L*_*base*_, with *L*_*Consistency*_, where the latter is scaled by a factor of 0.1:

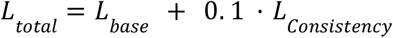

We adhered to the same learning rate schedule, perturbation parameters, batch size, and stopping criteria as delineated in the section discussing runtime experiments. All Visium data was preprocessed following the methodology outlined in the preceding section.

When comparing with STAlign we use the same default parameters outlined in their article^15^.

### 3D modelling

During the training process, for each individual sample *X*_*i*_ drawn from the complete stack *X*_1_, …, *X*_*N*_, where *N* is the total number of sections, a random reference point *X*_*reference*_ is chosen from the z-stack. Furthermore, an additional sample, *X*_*j*_, is selected at random from within the range *X*_*i*−3_ to *X*_*i*+3_, with a certain amount of noise introduced. Landmarks are identified for each sample in this triplet: *X*_*reference*_, *X*_*i*_ and *X*_*j*_.

The non-distorted *X*_*i*_ is registered to *X*_*reference*_ using both a rigid transformation (utilizing the Kabsch-Umeyama algorithm) and a TPS transformation, resulting in two different versions of the registered image, denoted as 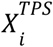 and 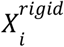. The noisy variant *X*_*j*_ is registered to the reference point using TPS, referred to as 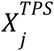.

Subsequent to registration, we compute the change in area, *dA*, before and after registration with TPS between *X*_*i*_ and 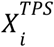. The loss function is then given as:

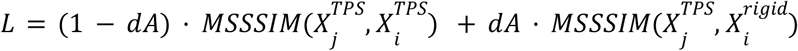

In this loss function, the first part is the product of MS-SSIM calculated between 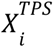 and 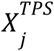, and (1 − *dA*). The second part is the product of MS-SSIM calculated between 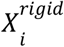 and 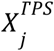, multiplied by *dA*.

This means that if the area changes significantly after registration, 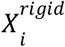 contributes more to the loss function, which could lead to a less optimal fit. This strategy compels the landmarks to function more as anchor points, ensuring increased stability throughout the z-stack.

We trained the model for 80 epochs, a batch size of 258 (the whole z-stack), resulting in 300 iterations per epoch, and with elastic noise with a sigma value of 5.5. The learning rate employed was 1e-4, using a learning rate scheduler with a step size of 10 epochs and a learning rate decay of 0.95.

All benchmark metrics were performed with the code from K. Kartasalo et al.^16^.

### Performance Evaluation on Multimodal Data

In the same way as in previous sections, we calculate a base loss by identifying landmarks and registering a sample with a noisy variant of itself. However, when we deal with multimodal data alignment, each data modality presents unique characteristics, distributions, and scales. This uniqueness can complicate direct comparisons using methods such as MS-SSIM, rendering them less meaningful. Therefore, for measuring the quality of alignment in this context, we need to employ an alternative proxy, distinct from MS-SSIM.

In our multimodal consistency loss computation, we employ the latent representations derived from the landmark detector. When aligning a sample from modality A with a sample from modality B, we run both samples through the landmark detector and obtain the activations of the first layer, represented as *Z*_*A*_ and *Z*_*B*_. We then gauge their similarity using the Root Mean Squared Error (RMSE), termed as the inter-consistency loss:

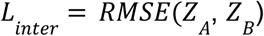

Moreover, we calculate intra-modality consistency by aligning samples within the same modality and leveraging MS-SSIM for loss computation. This intra-modality consistency mirrors the consistency loss outlined in the previous sections, where one section is registered to another within the same modality:

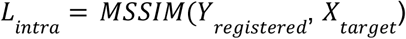

Analogous to prior sections, our base loss involves registering a noisy variant of a sample with another perturbed version of the same sample:

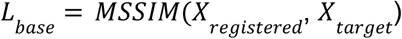

The final cost function amalgamates the base loss, inter-consistency loss, and intra-consistency loss. These are scaled by factors of 10 and 0.1, respectively, as determined empirically:

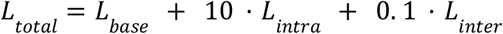

We followed the same protocol for the learning rate schedule, perturbation parameters, batch size, and stopping criteria, as detailed in the section regarding runtime experiments. As for Visium data, we maintained the same preprocessing steps described in the earlier section.

## Acknowledgment

This project has received funding from the European Research Council (ERC) under the European Union’s Horizon 2020 research and innovation program (grant agreement no. 101021019). The Erling Persson Family Foundation, the Swedish Cancer Society, and Swedish Research Council also supported the study.

